# Multiplexed temporally focused light shaping through a GRIN lens for precise in-depth optogenetic photostimulation

**DOI:** 10.1101/515908

**Authors:** Nicolò Accanto, I-Wen Chen, Emiliano Ronzitti, Clément Molinier, Christophe Tourain, Eirini Papagiakoumou, Valentina Emiliani

## Abstract

In the past 10 years, the use of light has become irreplaceable for the optogenetic study and control of neurons and neural circuits. Optical techniques are however limited by scattering and can only see through a depth of few hundreds µm in living tissues. GRIN lens based micro-endoscopes represent a powerful solution to reach deeper regions. In this work we demonstrate that cutting edge optical methods for the precise photostimulation of multiple neurons in three dimensions can be performed through a GRIN lens. By spatio-temporally shaping a laser beam in the two-photon regime we project several tens of targets, spatially confined to the size of a single cell, in a volume of 150×150×400 μm^3^. We then apply such concept to the optogenetic stimulation of multiple neurons simultaneously *in vivo* in mice. Our work paves the way for an all-optical investigation of neural circuits at previously unattainable depths.

## Introduction

Understanding communication among neurons and how they coordinate and integrate multiple signals is essential for discovering how neural circuits determine brain function and dysfunction. With a still growing toolbox of optogenetic reporters ^1,2^ and actuators ^3,4^, and the parallel development of advanced optical techniques, it is today possible to activate and inhibit neuronal activity with light while optically recording the evoked response. Ultimately this will enable all-optical neural circuit interrogation with single-cell and single-action potential precision, even in deep brain regions^5^.

One-photon (1P) wide field illumination using single optical fibers, enables the simultaneous illumination of large brain regions up to few mm deep and has already permitted to establish the links between neural activity and behaviour in different areas^6–11^. This light-delivery approach however lacks spatial selectivity. More sophisticated multi-point photonic probes^12–14^ achieved selective excitation of single brain layers but still lacked cellular resolution and did not permit concurrent optical recording of functional activity. By using 1P-computer generated holography (CGH) for patterned illumination through a fiber bundle coupled with a small objective, our group previously demonstrated single- and multi-neuron activation together with functional calcium imaging in freely moving mice^15^. However, the large dimension of the micro-objective (<2.5 mm diameter) limited this approach to superficial brain layers and the use of the fiber bundle did not preserve the holographic phase, thus constraining multi-spot generation to a single plane.

Two-photon (2P) excitation ^16,17^ combined with wave front shaping and opsin engineering allows millisecond manipulation of brain circuits with single cell resolution at multiple planes in three dimensions (3Ds). This was done either by generating multiple diffraction-limited spots that were scanned simultaneously across multiple cell somata^18–20^, or by using computer generate holography (CGH) to produce light patterns covering multiple cell somata at once^21^, thus optimizing the temporal precision of the photostimulation^22^. Recently, several research groups^23–27^ have shown that using a two-step wave front shaping combined with temporal focusing (TF)^28–31^ it is possible to generate multiple high resolution extended light patterns in 3D, a technique we named multiplexed temporally focused light shaping (MTF-LS). These approaches led to the first demonstrations of neural circuit manipulation in 3D^25,26^, yet the need of using conventional high numerical aperture (NA) objectives limited their use to circuits in superficial (≤ 300 μm) cortical areas or to *in-vitro* applications.

Micro-endoscopes (MEs), i.e. small optical probes that can be inserted into living tissues, represent a promising solution to extend optical brain manipulation to deeper brain structures. Most MEs are based on the use of gradient index (GRIN) lenses, which are small cylinders of glass of diameter < 1 mm and length of several centimetres, characterized by a gradual variation of refractive index in the radial direction^32^. In the last 15 years, GRIN lenses were studied and optimized for the 2P imaging of living tissues, e.g. implanted till a few mm deep inside the rodent brain^33–41^. *In vivo* functional calcium imaging was demonstrated through GRIN lenses both in 1P^42^ and in 2P excitation^43–46^. In Ref.^43^, 2P CGH was also performed through a GRIN lens (NA=0.5, diameter 0.5 mm) and a GRIN objective (NA=0.8, diameter 1 mm). Multiple diffraction-limited spots allowed the activity from several neurons to be imaged simultaneously through the GRIN lens. Extended light patterns were also generated through the GRIN objective (10 μm spots with axial resolution of ~ 22 μm). However, CGH was limited to a single plane and extended holographic patterns with single cell axial resolution were only shown through the larger and more invasive GRIN objective. Moreover, no experiment showed so far optogenetic photostimulation through GRIN lenses.

In this work we demonstrate an optical system combining our recent MTF-LS technique^27^ with the use of a GRIN lens (NA=0.5, diameter 0.5 mm), and we show that a small ME is suitable for the generation of multiple axially-confined 2P excitation spots matching target somata in 3D. Successively, using this system we demonstrate all-optical control of single or multiple neurons through the ME, by performing *in vivo* 2P optogenetic stimulation of target cells and reading out their induced activity with 2P calcium imaging.

## Results

### Optical setup

The optical setup coupling the MTF-LS^27^ to a GRIN lens ME is shown in Fig.1. In the optical characterization experiments we used a femtosecond fiber laser (Fidelity 10, Coherent), emitting at 1040 nm. The MTF-LS setup consisted of (1) a first SLM that determined the size and shape of the spot(s) to be generated at the sample plane; (2) a diffraction grating for TF; (3) a second SLM that multiplexed the axially confined spot(s) at arbitrary positions in 3D. The two SLMs were also used to correct for the aberrations of the system, in particular of the GRIN lens. To do so we maximized the 2P signal from a diffraction-limited spot made at the centre of the field of view (FOV) by introducing the appropriate first orders Zernike aberration corrections. We used an air objective (Olympus, UPLFLN 10X) with NA of 0.3 to focus into the GRIN lens, which we purchased from GRINTECH GmbH (model NEM-050-25-10-860-DM). It had a diameter of 0.5 mm and a total length of 9.86 mm, with NA of 0.19 on the objective side and of 0.5 on the sample side, resulting in a total field of view (FOV) of ~ 150 μm diameter^46^. For the optical characterization, the spots produced using MTF-LS through the GRIN lens were focused onto a thin rhodamine layer and imaged with a second objective (40x-NA 0.8 objective LUM PLAN FI/IR, Olympus) in transmission on a CCD camera (CoolSNAP HQ2, Photometrics). To characterize the axial resolutions, the rhodamine sample was vertically scanned together with the imaging objective with a piezoelectric z scanner (PI N-725.2A PIFOC). As discussed in Ref.^27^, the MTF-LS technique is compatible with several different beam-shaping strategies that one can perform with the first SLM. In the following we show the results of (i) multiplexed temporally focused computer generated holography (MTF-CGH) and (ii) multiplexed temporally focused multi shapes (MTF-MS) through the GRIN lens.

**Fig.1.**
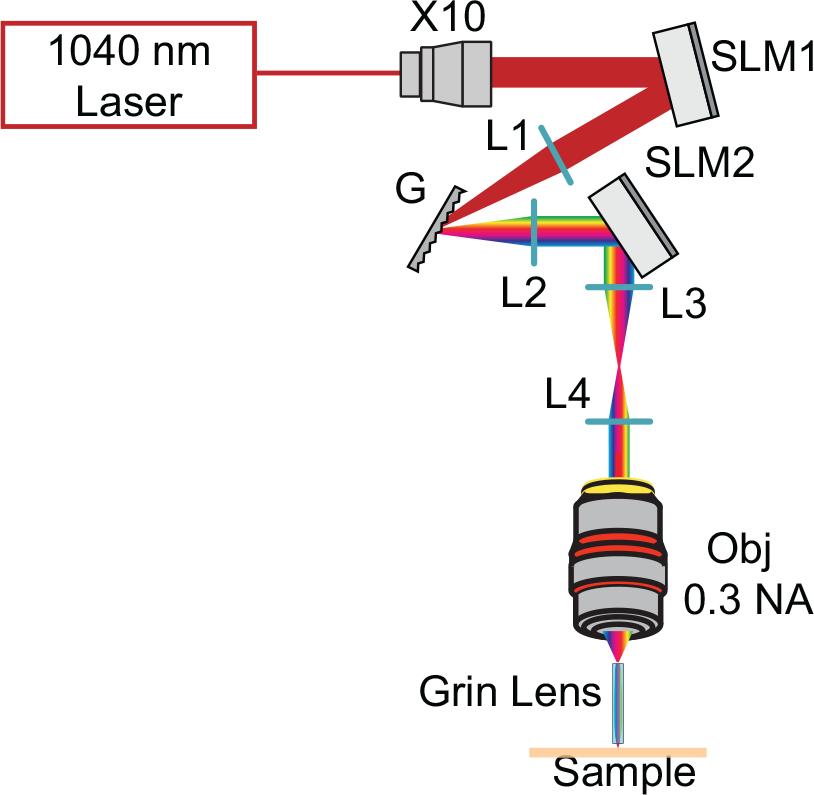
Schematics of the optical setup used in the experiment comprising a 10 times beam expander, two SLMs, the diffraction grating (G) for TF, the appropriate lenses (L), the 10X, 0.3 NA air objective and the GRIN lens. For optical characterization the sample was a thin layer of rhodamine that was scanned in *z* for axial resolution measurements.

### MTF-CGH through the GRIN lens

In a first experiment we used the first SLM (see Fig.1) to generate a simple holographic shape (a 15-μm diameter round spot) that was focused onto the diffraction grating for TF. The second SLM multiplexed the axially confined spot at arbitrary positions in 3D. In Fig. 2a we compare the axial resolution of a 15-μm spot generated through the GRIN lens at the centre of the FOV to the case of a non temporally focused spot of the same size (red) and to that of a spot generated through the 0.3 NA air objective alone (with no GRIN lens) with TF (blue). As the green curve clearly suggests, when TF was not used, the axial resolution, calculated as the full width at half maximum (FWHM) of the curve, was excessively large (~ 80 μm), as a consequence of the relatively small NA of the GRIN lens. When using TF, the axial resolution at the centre of the FOV improved to ~ 20 μm.

**Fig. 2.**
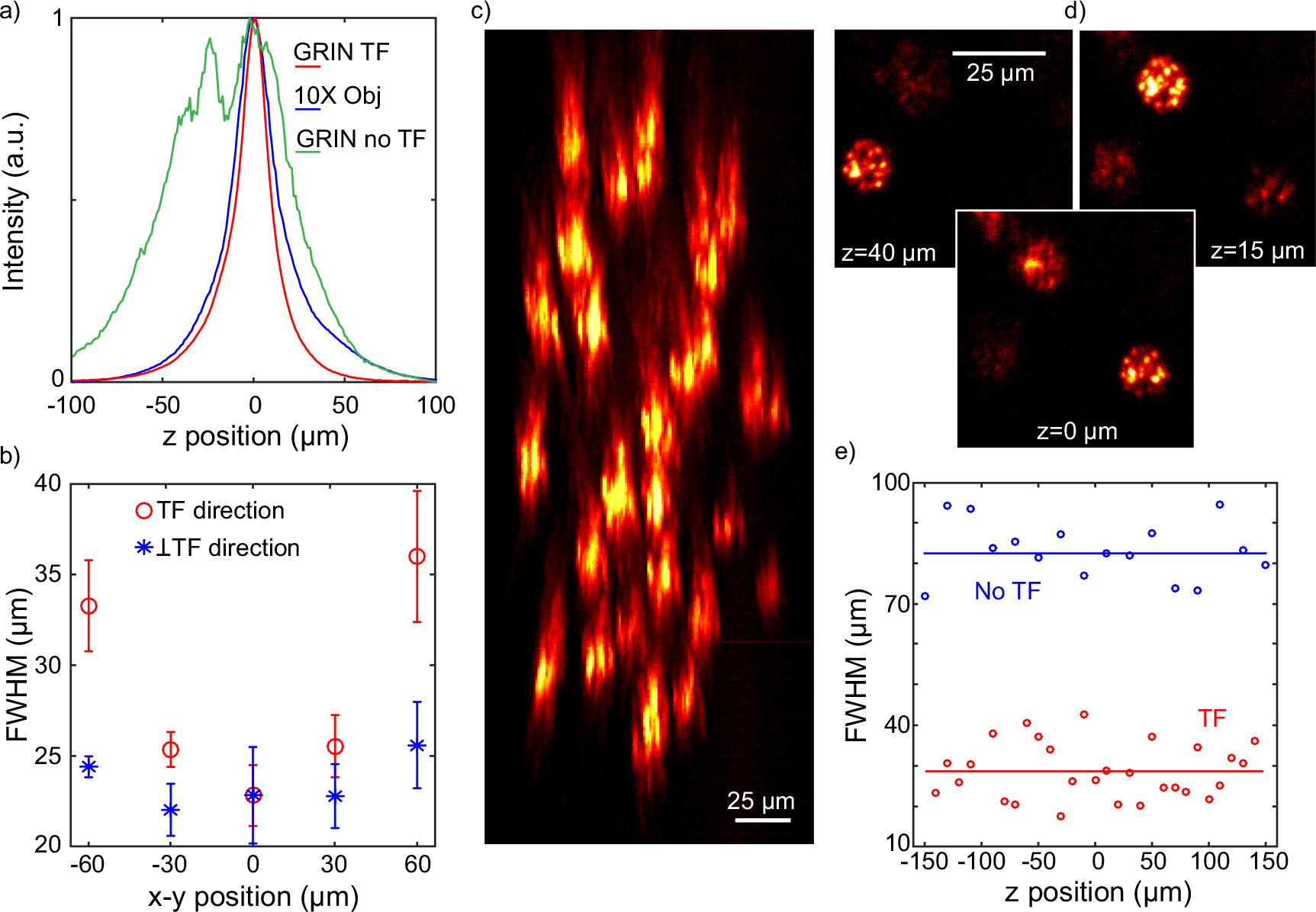
Results obtained using the MTF-CGH technique though the GRIN lens. a) Axial profiles for a 15 μm holographic spot at the center of the FOV when using the GRIN lens with TF, without TF and the 10X objective alone with TF. b) Behavior of the FWHM of one holographic spot displaced along the TF (*x*) or the perpendicular (*y*) direction. The data are the average over 6 different repetition of the same experiment and are given with an error bar calculated as the standard deviation over all the repetitions. c) Lateral view of 28 holographic spots distributed in 3D. d) In plane view of 3 holographic spots at 3 different depths. e) Comparison of the axial resolutions with TF (28 spots) and without TF (16 spots).

In Fig. 2b we compare how the axial resolution of the holographic spots changed as a function of the lateral displacement both in the *x* (direction of dispersion of the diffraction grating) and in the *y* (perpendicular to the dispersion) direction (Fig. 2b). From the graph one sees that the axial resolution worsened in the TF direction (i.e. the *x* direction) when moving away from the centre much faster than for the *y* direction. This asymmetry, as detailed in the discussion, was not observed in previous experiments ^27^, and could be due to a loss of colours through the GRIN lens when displacing the holographic spot in the TF direction. In any case, the temporally focused spots maintained an axial resolution better than 25 μm across 120 μm in the *y* direction and across 60 μm in the *x* direction, reaching the value of 35 μm at 60 μm from the centre along *x*.

We next used the MTF-CGH system to produce 28 holographic spots in a FOV of 150×150×400 μm^3^ through the GRIN lens. Fig. 2c,d show respectively a lateral projection (*xz* plane) of the spots and the *xy* view at three different *z* positions, illustrating how different spots are focused at different depths. The plot in Fig. 2e compares the axial resolutions of the 28 temporally focused spots with a similar experiment in which we generated a distribution of 16 spots without using TF. The difference between the two cases is striking; the mean axial resolution was 28±7 μm in the former and 82±7 μm in the latter case (the error was calculated as the standard deviation over all the experimental repetitions). From these results one can clearly see that advanced optical methods, here CGH and TF, can be implemented through a GRIN lens ME. Moreover, TF is essential to preserve single cell axial resolution when using low-NA GRIN lenses to create extended 2P excitation spots.

### MTF-MS through the GRIN lens

As described in Ref.^27^, the MTF-MS method relies on a mixed phase/amplitude shaping approach, capable of simultaneously producing different speckle-free shapes, with an overall better axial resolution with respect to MTF-CGH, and to independently multiplex them at the sample plane. Here we used MTF-MS in two different configurations (Fig. 3): in the first one we generated a mixture of round and stars (for a total of 20 spots) (Fig. 3a,b); in the second one we generated 16 identical round spots (Fig. 3c). In Fig. 3a we show the lateral view (*xz* plane) corresponding to the case of two different shapes, whereas in Fig. 3b and c we show the projections along the *z* direction in the case of two and one shapes respectively. As expected, in both cases, we could generate a 3D distribution of shapes with improved axial resolution and intensity homogeneity with respect to Fig. 2. As summarized in Fig. 3d, the axial resolution was around 22±6 μm in the case of a single shape and 26±7 μm when using two different shapes. As discussed later, contrary to previous results through a normal objective^27^, when using the GRIN lens we could not easily generate more than two shapes, probably due to the larger aberrations of the system.

**Fig. 3.**
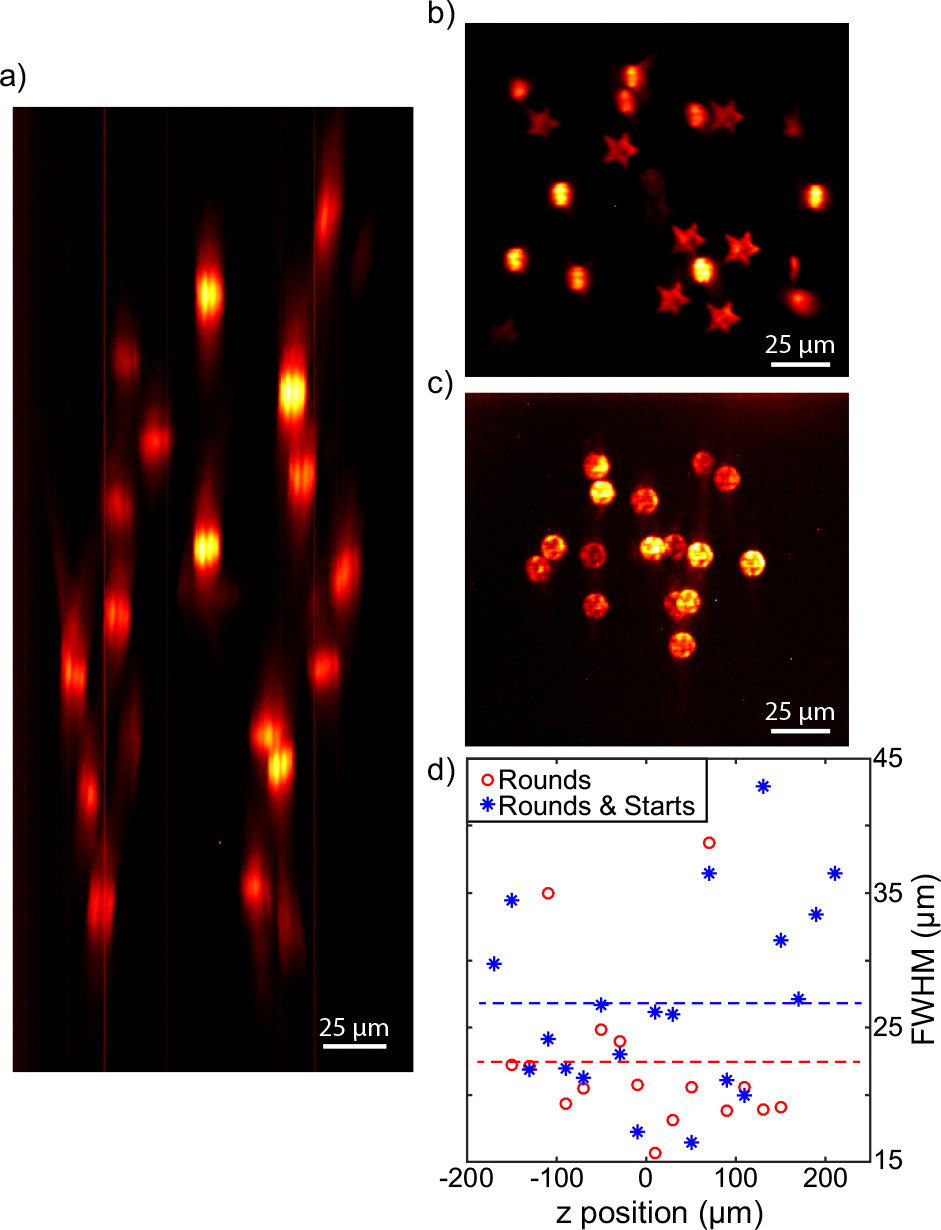
Results obtained using the MTF-MS technique though the GRIN lens. a) Lateral view of 20 spots distributed in 3D when using the first SLM to produce two shapes. b, c) Projections along the *z* direction in the case of two (rounds and stars) and one (rounds) shapes produced by the first SLM respectively. e) Comparison of the axial resolutions for the two cases.

### In vivo setup

We next used the GRIN lens based ME to perform concurrent holographic stimulation of single or multiple cells and optical readout of population activity *in vivo* in mice. As previously reported, a subset of neurons can be reliably co-labelled with opsins and calcium sensors by infecting cortical neurons of GCaMP6s transgenic mouse line GP4.3^47,48^ with ReaChR^49^ viral vectors ^50^.

In the *in vivo* experiments, we mounted the GRIN lens ME on a microscope already equipped for 2P optogenetic stimulation and 2P scanning imaging, described in Ref.^50^. In this case, we used a more basic version of the holographic setup shown in Fig. 1, comprising only one SLM and a diffraction grating, which could therefore only generate excitation spots on a single plane. The laser sources were an amplified fiber laser (Satsuma HP, Amplitude Systemes) emitting at 1030 nm for the photostimulation and a tuneable Ti-Sapphire laser (Coherent Chameleon Vision II) for imaging. The imaging laser beam was raster scanned on the sample with *xy* galvanometric mirrors (6215H series, Cambridge Technology) and the emitted fluorescence was collected back through the GRIN lens and objective, separated from the excitation path with a dichroic mirror, and directed to two photomultiplier tubes.

After viral infection of ReaChR at the depths of ~250 μm and ~500 μm in the mouse primary visual cortex (V1), we performed acute craniotomy and removed the dura mater to expose the brain in anesthetized mice. All the *in vivo* experiments were performed in the anesthetized mouse V1 using the GRIN lens to image and photo stimulate through the craniotomy. By moving the objective and the GRIN lens together with a *z* motor we could change the focus of the system from the brain surface down to ~250 μm deep (when the GRIN lens was in contact with the brain surface), possibly deeper if the GRIN lens slightly penetrated the brain.

### All-optical control of neuronal activity through GRIN lens *in vivo*

Through the GRIN lens, we were able to resolve on average 34±3 individual cortical neurons in a FOV of 150×150 μm^2^ (mean±s.e.m., 9 FOVs). As Fig. 4a shows, we observed GCaMP6s fluorescence changes in single (5 FOVs) or multiple (5.2±0.7 cell, 5 FOVs) target neurons by stimulating with 10 light-pulses at a repetition rate of 11.84 Hz, through 12-μm diameter circular holographic spots at a threshold illumination intensity (defined as the intensity resulting in ≥0.5 activation probability in target cells, see Methods) of 0.39±0.09 mW/μm^2^. The imaging power was kept at 90±14 mW, using a scanning speed of 5.92 or 11.84 Hz. Higher imaging powers led to an increased GCaMP6s fluorescence in cytosol, probably due to the slow channel kinetics of ReaChR^47^.

**Figure 4.**
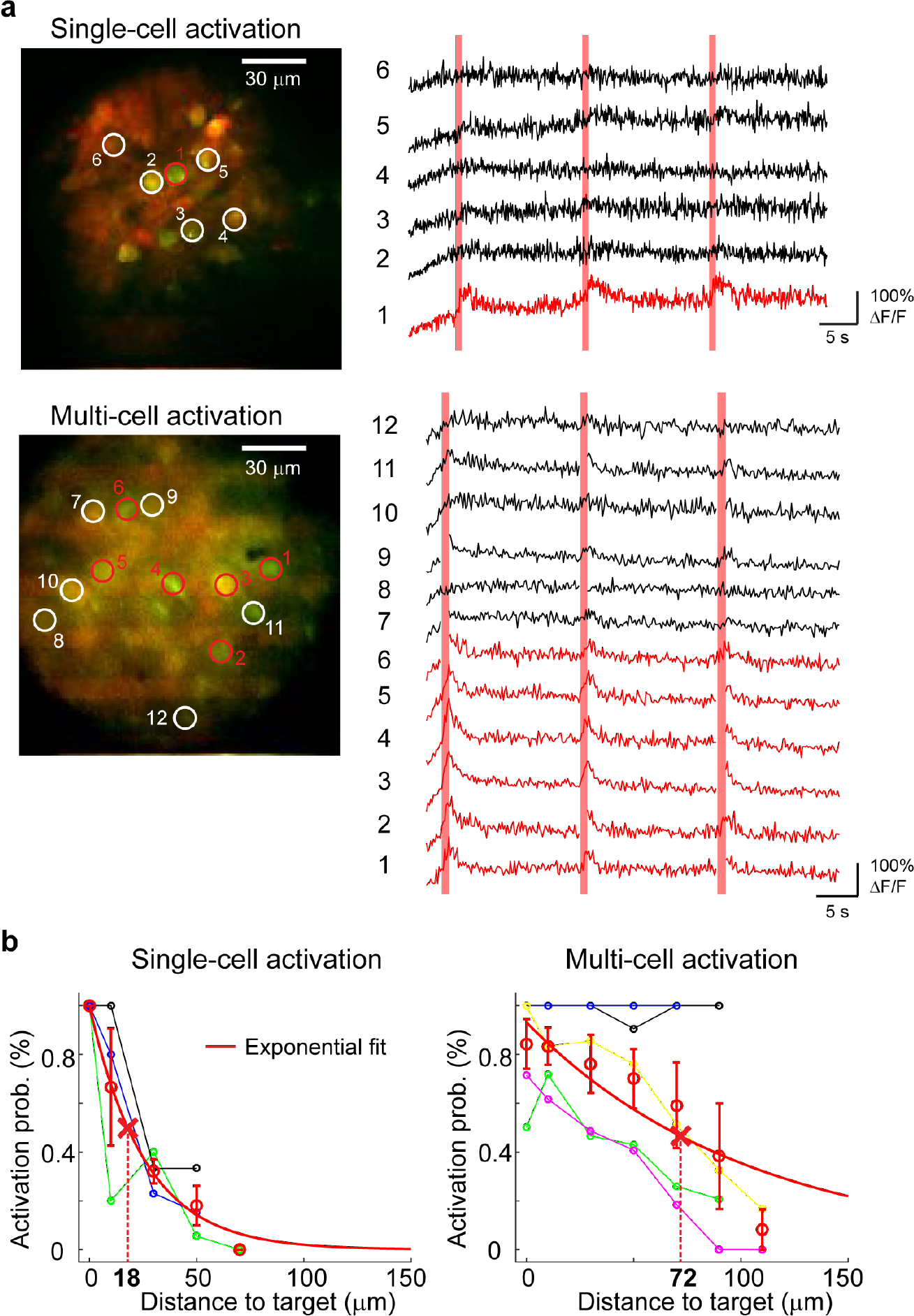
All-optical control through the GRIN lens in mouse V1 in vivo. (a) Upper panels show a case of single-cell activation experiment at the depth of ~110 μm below the brain surface. The red circle, in the average projection profile of 2P image intensity from 2 channels, represents the target cell and the 5 white circles are examples of non-target cells. The corresponding calcium traces are displayed at the right, with red vertical bars denoting the photostimulation epochs. Lower panels show a case of multi-cell activation experiment with 6 target cells (red circles) and 6 examples of non-target cells (with circles) at the depth of ~250 μm. Of note, the consecutive lines of bright pixels in the 2P images (upper and lower left) are photostimulation artefacts. Calcium signal in imaging frames from ROIs that were affected by stimulation artefacts is not shown. (b) Color lines (except red) indicate activation probability of target and non-target cells in relation to the distance to target cell for 5 FOV of single-cell activation (left) and 5 FOV of multi-cell activation (right). Red circles with error bars as mean±s.e.m. Red solid lines represent the exponential fits of activation probability. Dash lines indicate the distance where non-target cells displayed 50% target cells’ activation probability at the threshold illumination intensity. The distance between non-target and target cell is distributed in 20-μm bins.

To examine the spatial selectivity of photostimulation, we determined the relationship between the activation probability (see methods) of non-target cells and their distance to the target cells ^50^. We found the critical distance where non-target cells displayed 50% of target cells’ activation probability at the threshold stimulation intensity (see methods) to be 18 μm for single-cell activation (4 FOVs) and as 72 μm for multi-cell activation (5 FOVs) (Fig. 4b). Of note, for 2 FOVs in multi-cell activation experiments, non-target cells displayed >0.9 activation probability at the threshold stimulation intensity up to 100 μm away from a target cell, which implied neuronal network activation, thus reducing the spatial selectivity of photoactivation.

In sum, we demonstrated in vivo all-optical control of a single and multiple neurons through a GRIN lens. Out-of-target activation could result from a combination of neurite activation, smaller FOV and the relative low NA of the GRIN lens.

## Discussion

In this work we developed a ME, based on the combination of MTF-LS with a GRIN lens, for the 2P excitation of multiple targets in 3D and we combined it with functional imaging to enable cell-targeted all-optical interrogations of deep brain regions *in vivo*.

Using the TF technique and holographic multiplexing we could generate 3D distributions of extended excitation spots over a total FOV of ~ 150×150×400 μm^3^, keeping at the same time an axial resolution better than 30 μm. Importantly, our results demonstrate that the TF effect is maintained through the GRIN lens, despite its larger aberrations^51^. As shown in Fig. 2, this is crucial to keep a good axial confinement when generating large excitation spots, as the relatively small NA of the GRIN lens (0.5 on the sample side) would lead to a 4 times worse axial confinement if TF were not used therefore preventing targeted excitation. We showed that the GRIN lens is compatible with different MTF-LS techniques, such as MTF-MS. This latter configuration has the advantage of generating uniform speckle-free shapes with better axial resolution and gives the flexibility to separately multiplex different shapes. The increased light uniformity could be advantageous for parallel multi target imaging, while the possibility of generating distinct shapes could be used in experiments requiring activating different cellular compartments or different neuronal populations with variable cell size.

Compared to a MTF-LS conventional microscope, the use of a GRIN lens imposes few limitations (Fig. 2 and 3). First, we observed an asymmetric deterioration of the axial resolution when moving holographic spots in the x,y plane. Precisely, the axial resolution worsened more in the TF (*x*) than in its perpendicular direction (*y*) (Fig. 2b), a feature we did not observe when using a conventional high NA objective^27^. One can think at the GRIN lens as a doublet, which collimates the light it receives at the input and focuses it back at the output. There is therefore inside the GRIN lens an effective Fourier plane, namely a plane at which the wavelength components (or colours) of the laser pulse are separated in space. Shifting the spots at the entrance of the GRIN lens results in some portion of the beam (at some intermediate plane inside the GRIN lens) being closer to the edges of the GRIN lens, which are more prone to effective loss of light and aberrations. Intuitively, one can think that moving the spots in the *x* direction away from the centre, results in some of the wavelength components at an intermediate plane inside the GRIN lens being closer to the edge of the GRIN lens, producing a net loss of colours and hence of axial resolution. Moving the spots in the *y* direction instead, causes a similar loss of light for all the colours, leading to a net decrease of light, but not of axial resolution.

The second limitation was in the maximum number of different shapes we could create when using MTF-MS. While using a conventional high NA objective allowed us to simultaneously generate up to 4 different shapes^27^, using the GRIN lens we could only generate 2 different shapes. MTF-LS requires illuminating different portions of the second SLM with each shape, which in turns means entering into the microscope objective at different positions of its back entrance. This results in a tilted propagation at the sample plane for those shapes entering in the objective away from the centre, as also explained in^23^. If this problem had negligible consequences in the case of a high NA objective^27^, when using the GRIN lens it resulted in a loss of axial resolution and spot quality as the number of different shapes created by the first SLM increased. This was probably due to the larger aberrations of the GRIN lens compared to a normal objective^51^. A deeper study of the aberrations, including a method to use the two SLMs to correct for aberrations at multiple planes and/or lateral positions (and not only at the centre of the FOV as demonstrated here), or the future availability of optimized GRIN lenses with lower aberrations could help to overcome these limitations.

Low-NA GRIN lenses are currently the preferential choice for in-depth functional imaging in living animals as they permit minimal tissue damage compared to larger-diameter high-NA versions. We demonstrated the possibility to couple them with patterned illumination and temporal focusing enabling all-optical neuronal investigations *in vivo*. We were able to evoke calcium transients via soma illumination in cells co-expressing GCaMP6s and ReaChR in the mouse visual cortex *in vivo*. For single-cell activation through the GRIN lens, we found a critical distance of 18 μm, where non-target cells displayed activation probability that is 50% of that in target cells. This distance significantly increased (nearly 4 times) when multiple cells were simultaneously stimulated, as expected from the combined use of a non-somatic opsin^52,53^ and a relatively low NA GRIN lens and resulted from indirect synaptic activation. Out of target activation can thus be prevented by using somatic opsins and possibly GRIN lenses with a bigger FOV or by further improving the axial resolution, e.g. by using higher NA GRIN lenses.

In the present study, we have limited the demonstration of all-optical neuronal control to a single-plane. The combination of MTF-LS with several volumetric imaging implementations as upstream divergence control^20,21^ remote focusing^54^ or extended depth-of-field scan^55^ will enable to extend these demonstration to multiple planes. The use of chronic implantation of GRIN lenses (as shown for example in^43,46^) will also enable to extend this approach to the optical manipulation of deeper brain regions.

## Methods

### Animal preparations

All animal experiments were performed in accordance with the Directive 2010/63/EU of the European Parliament and of the Council of 22 September 2010. The protocols were approved by the Paris Descartes Ethics Committee for Animal Research with the registered number CEEA34.EV.118.12. Mice were anesthetized with intraperitoneal injection of a ketamine-xylazine mixture (0.1 mg ketamine and 0.01 mg xylazine/g body weight) during viral injection and with isoflurane (2% for induction and 0.5-1% for experiment) during photostimulation experiments. To express both opsin and calcium indicator in the same neuronal sub-population, stereotaxic injections of the viral vector AAV1-EF1α-ReaChR-dTomato were performed in 4-week-old male or female mice of transgenic line GP4.3 (The Jackson Laboratory), which constitutively expresses the calcium indicator GCaMP6s ^48^. Viral vectors of 1-1.5 μL were infused at either 250 μm or in combination with 500 μm to target cortical neurons in the right primary visual cortex at a speed of 80-100 nL/min. Acute experiments of holographic stimulation were performed 6-12 weeks after viral infection.

### Holographic stimulation and calcium imaging through GRIN lens in vivo

The *in vivo* endoscope system was integrated to an existing custom-made 2P all-optical system thoroughly described in^50^. Specifically, a 0.5 NA GRIN lens was positioned by means of a custom-designed holder underneath a 10x microscope objective coupled with a SLM-based 2P patterned photostimulation and a 2P galvo-based scan imaging system. The GRIN lens holder was composed of kinematic mounts allowing X,Y,Z translation and rotation of the pitch and yaw axes of the GRIN lens, thus ensuring GRIN lens alignment to imaging and photostimulation paths. GRIN lens holder and objective were jointly connected to an axial motor allowing beam refocusing in the sample.

Initially, GRIN lens and objective were axially translated towards the head of the mouse till the image of brain surface appeared. That set the zero position of the GRIN lens, where its tip was at a distance of 250 μm (i.e., the optical system’s working distance) away from the brain surface. To perform all-optical experiments on a sub-population of cortical neurons co-expressing the opsin ReaChR and the calcium sensor GCaMP6s located up to 250 μm below the brain surface (i.e. layer 2/3), we axially refocused the GRIN lens/objective system like with an ordinary 2P objective microscope until at the brain surface.

For acute in vivo experiments, mice were anesthetized with isoflurane (as described above) and the skull was exposed after subcutaneous application of lidocaine. A custom-made head-plate was attached to the skull using dental cement (Unifast Trad, GC). A circular craniotomy of 2-3 mm diameter was made over the injection site and the dura mater was removed.

Two-photon imaging of GCaMP6s calcium signal and the fluorophore labelling of dTomato was carried out by exciting with a scan beam provided by a Ti:Sapphire laser emitting at 920 nm (Coherent Chameleon Vision II). SLM-based patterned photostimulation was provided by using high-energy fiber laser-pulses at 1030 nm (Satsuma HP, Amplitude Systemes).

Target cells were selected based on a high-resolution reference image (512×512 pixels) of the red channel collecting signal from ReaChR-dTomato. Through single or multiple circular holographic spots of 12-μm diameter, the target somata were simultaneously illuminated with 10 holographic light-pulses of 5 or 10 ms at 11.84 Hz. Concurrently, the neuronal population activity of GCaMP6s calcium signal in a 150×150 μm^2^ FOV was monitored by scanning at a frame rate of 5.92 or 11.84 Hz with 128×128 pixel resolution.

During all-optical experiments, the photostimulation laser induced artefactal excitation of fluorophores, which appeared as consecutive lines of bright pixels. The image frames of GCaMP6s signal from regions-of-interest (ROIs) that were affected by stimulation artefacts were removed in post-processing.

### Data analysis

Image analysis was performed by using ImageJ and MATLAB (Mathworks). ROIs covering individual target and non-target cell somata were manually selected in ImageJ based on both red (ReaChR-dTomato) and green (GCaMP6s) channels. The time-lapse fluorescent signal of GCaMP6s from all ROIs was exported in MATLAB. Image frames from ROIs that are affected by the photostimulation artefacts were removed in analysis. For each ROI, the relative percentage change of GCaMP6s fluorescence was computed as ΔF/F=(F-F_0_)/F_0_, where F_0_ was the average raw fluorescence signal 3-0.2 s before photostimulation started. A cell was considered activated if the average ΔF/F 1 s after the last illumination onset was significantly larger compared to that 1 s before the first illumination onset (right-tailed paired t-test with a significance level of 0.05). The relationship between activation probability and distance-to-target was determined similarly as previously described ^50^. Specifically, the threshold stimulation intensity was determined when the activation probability of target cells >=0.5. The average values of activation probability at different distance-to-target were described by fitting with an exponential function.

## Acknowledgments

We thank Vincent de Sars for software developing, Coherent Inc. for the loan of the Fidelity laser, This research received support from the ‘Agence Nationale de la Recherche’ (grant ANR-15-CE19-0001-01, 3DHoloPAc), the Human Frontiers Science Program (Grant RGP0015/2016), the Fondation Bettencourt Schueller (Prix Coups d’élan pour la recherché française), the Getty Lab, the National Institute of Health (Grant NIH 1UF1NS107574 - 01) and Axa research funding. N.A. received funding from the European Union’s Horizon 2020 research and innovation programme under the Marie Skłodowska-Curie grant agreement no. 746173. E.R. received funding from the European Research Council SYNERGY Grant scheme (HELMHOLTZ, ERC Grant Agreement # 610110). IWC received funding from the European Union’s Horizon 2020 research and innovation program under the Marie Skłodowska-Curie grant agreement no. 747598.

## Data availability

The datasets generated during and/or analysed during the current study are available from the corresponding author on reasonable request.

## Author contributions

N.A., IW.C. and E.R. designed the experiments. N.A. and C.M. built the 3D holographic micro-endoscope system, performed the optical experiments and analysed the optical data. E.P. participated in the development of the optical system. IW.C. performed animal surgery. E.R and E.P built the scanning imaging and 2D stimulation microscope. C.T. designed and built the GRIN lens holders to align the endoscope both for the optical and the in vivo setups. IW.C., E.R. and N.A. performed *in vivo* experiments. IW.C. analysed the in vivo data. N.A., IW.C. and V.E. wrote the manuscript with the contribution of all the authors. V.E. conceived and supervised the project.

